# bulkAnalyseR: An accessible, interactive pipeline for analysing and sharing bulk multi-modal sequencing data

**DOI:** 10.1101/2021.12.23.473982

**Authors:** Ilias Moutsopoulos, Eleanor C Williams, Irina I Mohorianu

**Affiliations:** Wellcome - MRC Cambridge Stem Cell Institute, Jeffrey Cheah Biomedical Centre, University of Cambridge, CB2 0AW, UK

**Keywords:** bulk sequencing, mRNAseq, ChIPseq, epigenetics sequencing, quality checking, differential expression, gene regulatory networks, multi-omics integration

## Abstract

Bulk sequencing experiments (single- and multi-omics) are essential for exploring wide-ranging biological questions. To facilitate interactive, exploratory tasks, coupled with the sharing of easily accessible information, we present **bulkAnalyseR**, a package integrating state-of-the-art approaches using an expression matrix as the starting point (pre-processing functions are available as part of the package). Static summary images are replaced with interactive panels illustrating quality-checking, differential expression analysis (with noise detection) and biological interpretation (enrichment analyses, identification expression patterns, followed by inference and comparison of regulatory interactions). **bulkAnalyseR** can handle different modalities, facilitating robust integration and comparison of cis-, trans- and customised regulatory networks.

**bulkAnalyseR** is available on CRAN and GitHub, with extensive documentation and usage examples (https://github.com/Core-Bioinformatics/bulkAnalyseR, https://cran.r-project.org/web/packages/bulkAnalyseR/)

**Contact:** Irina Mohorianu

## 1 Background

Bulk sequencing plays a key role in modern biomedical research, as a primary resource for generating hypotheses [1]. The wide diversity of available methods, each with specific trade-offs on speed, robustness, and reproducibility, hinders swift exploration of datasets. Options include bespoke analyses (through dedicated bioinformatics support), commercially available solutions (often expensive, and with limited flexibility for customisation) or fragmented pipeline-components relying on intermediary conversions of inputs/outputs [2, 3, 4].

Commonly-used tools/pipelines are rarely self-sufficient, i.e. cannot perform end-to-end analyses, and hence have limited appeal to researchers with restricted bioinformatics resources. *VIPER* [5] provides an end-to-end workflow starting from fastq files, but requires computing skills for setting up analyses and offers restricted scope for output customisation with user-controlled parameters (thresholds). *BioJupies* [6] is simpler to run and boasts a larger selection of outputs; however, it restricts the customisation of individual components e.g. to selecting genes; additionally, fewer options for the biological interpretation of outputs are available. Critically, neither offers support for complex differential expression (DE) tasks, beyond simple pairwise comparisons, limiting the biological interpretations of complex experimental designs. *Searchlight* [7] alleviates some of these issues, focusing on an interactive exploration of results; the inputs are tables of DE genes, relying on several *a priori* steps (QCs and DE inference). It also assumes expertise for setting up the environment and retrieving outputs; moreover, the resulting plots require additional tweaking to meet publication requirements.

Current state-of-the-art analyses go beyond generating lists of DE genes, tapping into causal (gene interaction) inference, employing single- or multi-omics inputs [8, 9]; however, most Gene Regulatory Network (GRN) inference tools are standalone. An early effort linking prediction and visualisation [10], underlines the lack of and need for direct, customisable bridges between analyses and biological interpretations. Visualisation tools such as *Cytoscape* [11] facilitate hypothesis generation, but require external definitions of the networks. Moreover, GRN inferences are often performed on the entire dataset, generating complex, difficult-to-interpret networks [12] and lacking in robustness due to intrinsic variability [13]. *GeNeCK* [14] attempts, in a web-based setting, to link GRN inference and interpretation, but is decoupled from other steps such as DE analysis.

**bulkAnalyseR** enables the analysis of single- and multi-omics bulk-sequencing data, in an interactive Shiny interface, with several customisable components, from noise detection [13] to the identification of patterns [15] and GRN prediction [4]; the latter accommodates cis-, trans- and custom interactions. The setup facilitates a seamless transition to publication-ready figures without additional bioinformatics support. The resulting web-app is easily publishable, incentivising open and reproducible research.

## 2 Results and Discussion

### bulkAnalyseR

provides an accessible, yet flexible end-to-end framework for analysing bulk-sequencing datasets, without relying on prior programming expertise (Fig. 1, Fig. S1). It generates a shareable Shiny app in two lines of code; all resulting plots and tables can be downloaded individually and the underlying code for generating the outputs can be easily reproduced.

**Figure 1:**
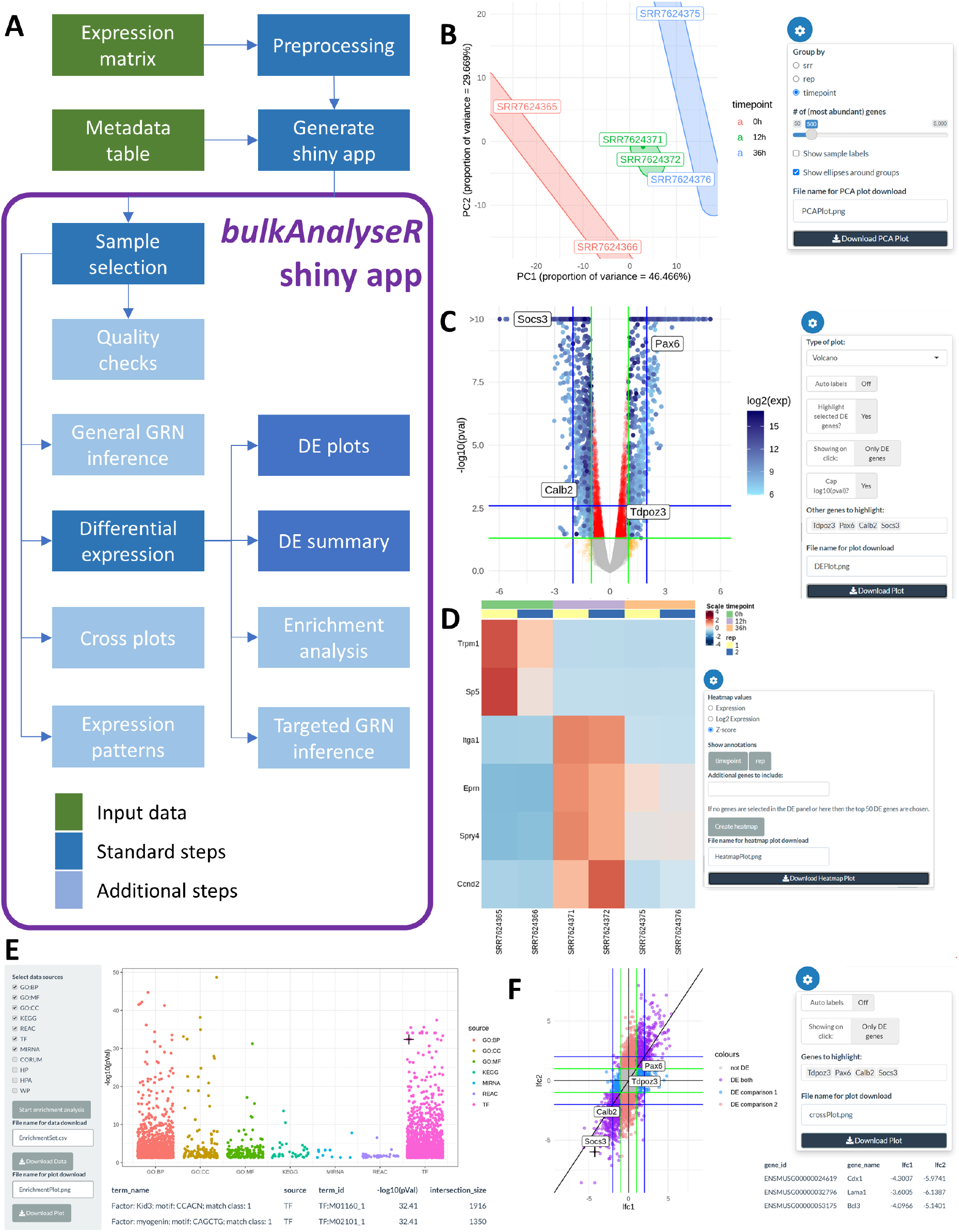
Workflow and examples of bulkAnalyseR summary plots on bulk mRNAseq data [8]. A) Workflow diagram of the main components of the pipeline and package; further details are presented in Fig. S1. B) PCA plot of all samples illustrating sample similarity. C) Volcano plot showcasing DE genes between the 0h and 12h samples. D) Heatmap of Z-scores across samples for the top differentially expressed genes. E) Enrichment analysis results for the DE genes (enriched terms ordered are by significance). F) Cross-plot comparing two DE outputs (*log*_2_FC from 0h vs 12h on the x-axis and and 0h vs 36h on the y-axis).

### Inputs and pre-processing

The inputs for **bulkAnalyseR** are an expression matrix and a metadata table. The former contains counts per genes across samples (e.g. generated from BAM alignments with featureCounts [16]). The metadata table contains additional experimental information and represents the starting point for the DE analyses/comparisons. The pre-processing step (`preprocessExpressionMatrix` function) handles the counts-based noise detection using noisyR [13], which outputs a denoised, un-normalised expression matrix (Fig. S2). Several options are available for normalisation [17]: quantile (default, [18]), per-total [19], TMM [2] and DESeq2 [3]. Integration with, or replacement by, other pre-processing steps is also possible, as any expression matrix can be passed onto the next step.

Next, `generateShinyApp` checks the compatibility of inputs, and the expression matrix (denoised, normalised, by default) and a Shiny app is created. The resulting app is standalone and shareable e.g. via online platforms like shinyapps.io (https://docs.rstudio.com/shinyapps.io/) [further details are provided in Supplementary 7]. This setup increases access and promotes reproducibility of bioinformatics analyses.

### Interactive Visualisation of differential expression

A single-omics instance comprises several tabs: [a] Quality Checks (QCs) include a Jaccard Similarity Index (JSI) heatmap and a PCA dimensionality reduction, with groups based on the metadata information (Fig. 1B, Fig. S4A,B). This enables a high-level overview of the similarity across samples, reflecting the experimental design. [b] The differential expression (DE) tab includes DE summaries across selected comparisons performed using edgeR [2] and DESeq2 [3] with customisable parameters (significance and fold-change thresholds are user-adjustable); the outputs are tables of DE genes (Fig. S4C). [c] The DE results can be visualised in the Volcano/MA and DE summary panels which promote an interactive exploration of the data (Fig. 1C, Fig. S4E,F,H). [d] The interpretation of the DE genes commences with the enrichment tab, which overviews a gene set enrichment analysis (GSEA) using g:profiler [20]. The GSEA is performed on GO terms, KEGG and Reactome pathway terms, and regulatory features (miRNAs [21] and transcription factors [22]), Fig. 1E. Further information is obtained by [e] identifying simplified patterns that may underline regulatory interactions (Fig. S3I) and [f] comparing DE lists in cross-plots (Fig. 1F, Fig. S4D).

The differences observed between frequently-used DE inference pipelines (either due to normalisation details or criteria for the DE call), pertinently discussed in Li et al [23] and Moutsopoulos et al [13], are assessed in Supplementary 3 on the Yang et al dataset. The variability in results is significantly reduced by addressing the (technical) noise [13]. For this case study, we highlight these differences by contrasting the DESeq2 and edgeR DE outputs, when the pipelines are run on the same inputs (noisy and denoised matrices are assessed, alongside different normalisation methods).

### Visualisation of Gene Regulatory Networks

Data exploration continues by inferring networks of regulatory interactions (GRN tab), using GENIE3 [4]. To allow the visualisation of changes in topology or strength of the interaction (in terms of co-expression) across different subsets of samples, up to 4 inferences can be performed simultaneously. Focus genes can be selected from the set of DE genes; to increase the robustness of this analysis, we recommend using up to 3 indirect interactions from the focus genes (Fig. 2A, Fig. S4J). An UpSet plot summarises the common vertices across the resulting networks.

**Figure 2:**
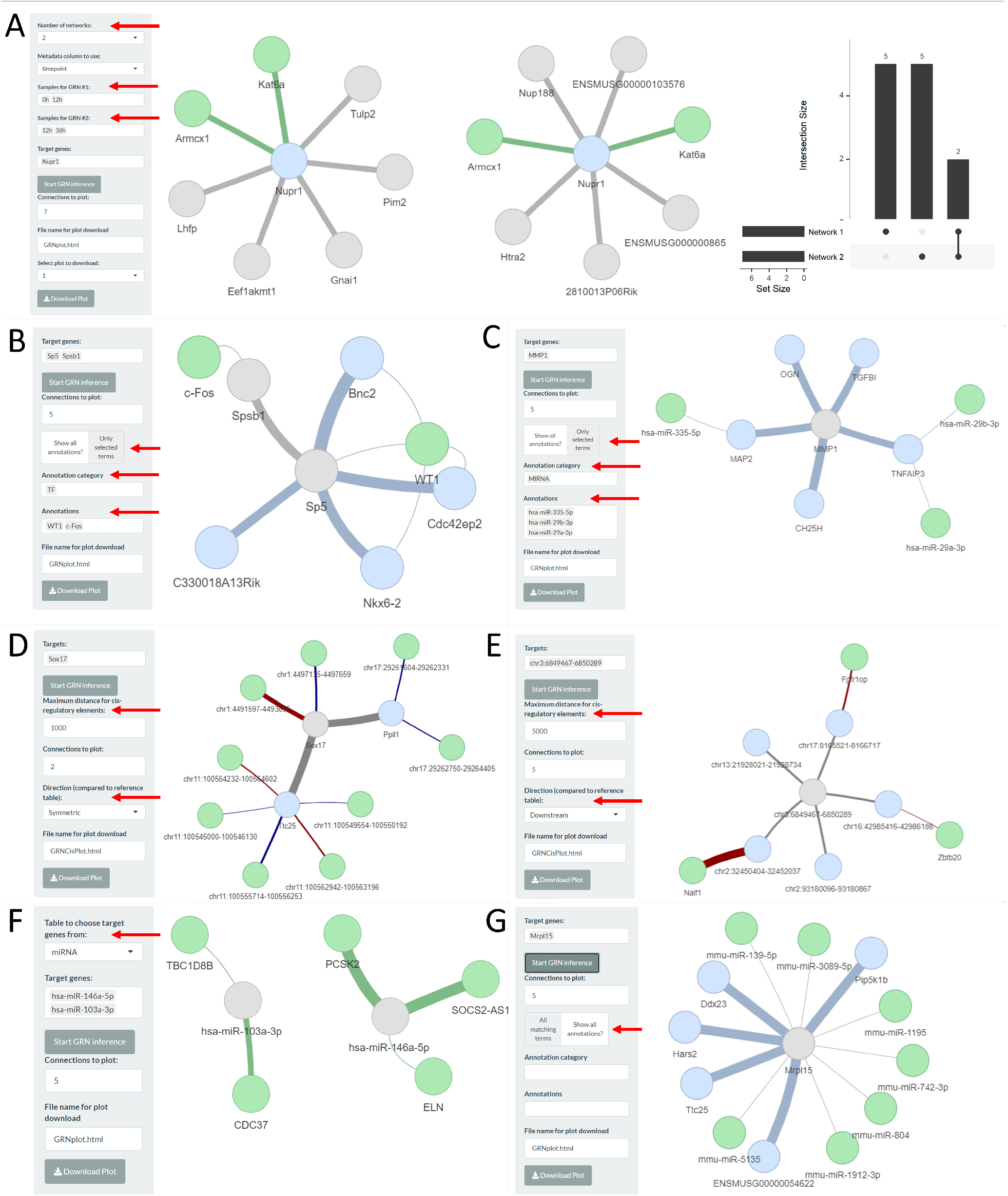
Examples of GRN analyses. A) Yang et al case study, visualisation of top 7 connections of gene *Nupr1*; the inference was performed on 0h vs 12h samples (left) and 12h vs 36h samples (middle). The width of the edges is proportional to the weight from the adjacency matrix. Target genes are shown in blue; common genes between the 2 networks are highlighted in green. The overlapping genes between the 2 networks are summarised in an UpSet plot (right). B) GRN inferred on 0h vs 12h DE genes augmented with enriched TF information (green, from g:profiler2). C) Li et al 2021 case study, GRN network inferred on control vs IDD DE genes for the mRNAseq modality, augmented with enriched microRNAs (green, from g:profiler2). D) Yang et al case study, GRN network inferred for *Ppil1* and *Ttc25* for 0h, 12h and 36h for the mRNAseq modality with cis-integration of the corresponding ChIpseq modality; in green the H3K4me3 ChIP peaks within 1kb of target genes are shown. Upstream peaks are highlighted with red edges; downstream peaks are highlighted with blue edges. E) GRN network inferred on the H3K4me3 modality, with cis-integration of genes (green) within 1kb downstream from the selected peaks. F) Li et al 2021 case stud, trans-integrated GRN, for mRNA and miRNA modalities; tthe GRN is miRNA-centric. G) Yang et al case study, GRN inferred on *Mrpl15* for 0h, 12h and 36h mRNAseq modality with customised miRTarBase integration (miRNAs linked to the selected mRNA are highlighted in green).

### Processing of other single modalitie

The input, based on an expression matrix, confers flexibility to **bulk-AnalyseR**. Other sequencing datasets, which can be summarised in an expression matrix, can be processed using this pipeline (we illustrate a case study of H3K4me3 ChIPseq data, Fig. S5, and of a small RNA dataset, with a focus on mature microRNA expression levels, Fig. S6). For these analyses, all steps except the GSEA, which cannot be defined on non-protein-coding genes, are available. The versatility of integrating multiple modalities or different experiments (of the same or mixed modalities) in the same framework is a promising first, interactive step to guiding mechanistic studies (e.g. multi-omics GRN inference).

### Multi-omics GRN integration bulkAnalyseR

also permits the integration of multiple modalities for inferring regulatory interactions (i.e. cis-, trans- and user-defined). Two inputs are required for the *cis-interactions*, which undergo soft/late integration: the focal modality and the non-focal modality, elements of which are located in proximity (e.g. 1kb, user-defined parameter), are used to augment the focal GRN. For example, the RNA-centric GRN can be augmented with information on cis-elements such as ChIPseq peaks (Fig. 2D); the two modalities are interchangeable (Fig. 2E). The proximity parameter can encompass entries from the non-focal dataset which are upstream, downstream or both, of the selected entries in the focal data. For the *trans-interactions*, the two inputs undergo complete/early integration, i.e. the two expression matrices, with identical columns/conditions, are concatenated. The GRN is inferred on the combined expression matrix; nodes are coloured according to the original source of the element (Fig. 2F). Given that the GRN inference is agnostic of expression amplitude, focusing just on variational patterns, no additional batch correction or over-normalisation is performed. The selected targets for the localised GRN can be selected from either input.

*Custom, user-defined interactions* can be added/ visualised; these rely on external information related to the vertices in the inferred GRN. Pre-loaded examples include adding miRNAs targeting protein-coding genes, retrieved from miRTarBase [21] (Fig. 2B) and interactions with transcription factors (Fig. 2C), retrieved from gprofiler2 [20]. Users can also embed their own customised interactions, supplied as a table (Fig. 2G). Example apps illustrating multi-omics GRN integration are [Yang et al, mRNA/ChIP integration](http://bioinf.stemcells.cam.ac.uk:3838/bulkAnalyseR/Yang2019_Integration/) and [Li et al mRNA/microRNA integration](http://bioinf.stemcells.cam.ac.uk:3838/bulkAnalyseR//Li2021/). A vignette detailing the integration of multi-omics and external data can be found [here](https://core-bioinformatics.github.io/bulkAnalyseR/articles/IntegratingMultiomics.html).

## 3 Conclusions

The aim of **bulkAnalyseR** is to enhance the interactive interrogation of single- and multi-omics data; additionally, the ability to share a stable instance of an analysis with the community has the potential to ease communication between research groups and generate new hypotheses that extend beyond the initial purpose of datasets. Moreover, the seamless integration of all steps in an end-to-end approach, from early quality control checks through to publication-ready figures, assists with data mining throughout the life cycle of an analysis.

More importantly, **bulkAnalyseR** provides the flexibility to integrate several modalities and datasets, and incorporate external databases through standard enrichment analysis, multi-omics integration and more customisable pipelines. This adaptability represents a valuable starting point towards more complex questions that may pave the road towards a better mechanistic understanding of the cause(s) behind the observed differences in expression.

## 4 Materials and Methods

### RNAseq modality

The single-omics mRNAseq functionality was exemplified using several mRNAseq samples from the *Yang et al*. [8] study (namely, 0h, 12h, and 36h post stem cell induction). The raw fastq files were downloaded from GEO (accession GSE117896). Initial quality checks were performed using fastQC (version 0.11.8) and summarized with multiQC (version 1.9) [24]. Alignment to the *M. musculus* genome was performed using STAR (version 2.7.0a) with default parameters [25]; the count matrices were generated using featureCounts (version 2.0.0) [16] against exon annotations from Ensembl (assembly GRCm38.p6).

### Epigenetic (ChIP) modality

The ChIPseq modality (for showcasing cis-interactions) was illustrated using several samples from the *Yang et al*. study; identical time points as for the mRNAseq samples were selected. Alignment to the *M. musculus* genome was performed with bowtie2 v2.4.2 (single-end mode, local mode default parameters) [26]; peak calling was performed using macs2 2.2.7.1 [27]. Peak aggregation across samples (for generating a unified expression matrix) was performed on the union of matched peaks. Peaks are matched if, per peak, its midpoint in one sample is within the peak genomic range in the other sample. Using pyBigWig 0.3.17 (deeptools, version 3.5.0 [28]), the corresponding amplitudes for these peaks were calculated (and scaled on the read length). The bigwig files were produced using bamCoverage (deeptools) with default parameters.

### Non-coding RNA modality

To showcase trans-regulatory interactions we focused on a mRNA/microRNA case study (*Li et al* [29]); the dataset describes intervertebral disc degeneration and comprises 3 control and 3 treatment (IDD) samples. The raw fastq files were downloaded from GEO (accession GSE167199). The mRNA samples were aligned to the *H. sapiens* reference genome using STAR (version 2.7.0a) with default parameters [25]; the count matrices were generated using featureCounts (version 2.0.0) [16] against exon annotations from Ensembl (assembly GRCh38.p13). Small RNA (microRNA) samples were aligned against *H. sapiens* mature microRNAs from miRBase [30] using PaTMaN [31] with 0 edits and 0 gaps. The mature microRNA expression was calculated as the number of exact copies of the miRNA sequence.

### Customised interactions

To illustrate customised interactions, we integrated the outputs from the enrichment step, and included the option to decorate a GRN with manually curated entries from miRTarBase [21], for post-transcriptional regulation, and TRANSFAC, [22], for transcriptional regulation.

### Differential expression and biological interpretation steps

For the case studies presented in the manuscript, the noise correction was performed using the counts approach from the noisyR package [13]. The normalisation was performed using (a) the preprocessCore package for quantile normalisation, (b) by scaling each value by one million divided by the reads in each sample for RPM, (c) the DESeq2 package for its internal normalisation, (d) the edgeR package for TMM, and (e) by dividing each value by the median expression in each sample for the median nomalisation.

The differential expression call is performed using the DESeq2 [3] or edgeR [2] pipelines, using default parameters (with data pre-normalised), on a comparison of the two conditions specified. The enrichment analysis is based on the gprofiler2 package [20] with a custom background of all genes expressed in the dataset. The gene regulatory network inference is performed using the GENIE3 package [4] focusing on user-defined regulatory targets. When more than one network is generated, edges appearing in more than one network are highlighted. An UpSet plot [32] summarises the intersections and specific differences of the GRNs.

## 5 Declarations

All authors read and approved the submitted version of the manuscript and declare no competing interests.

All authors acknowledge the constructive feedback from the Core Bioinformatics group and the support by the Core grant awarded to the Wellcome-MRC Cambridge Stem Cell Institute.

All authors contributed to the **bulkAnalyseR** package, and wrote and revised the manuscript.

## References

[1] Rory Stark, Marta Grzelak, and James Hadfield. RNA sequencing: the teenage years. Nat Rev Genetics, 20, 07 2019.

[2] Mark D. Robinson, Davis J McCarthy, and Gordon K Smyth. edgeR: a Bioconductor package for differential expression analysis of digital gene expression data. Bioinf, 26(1):139–140, 11 2009.

[3] Michael I Love, Wolfgang Huber, and Simon Anders. Moderated estimation of fold change and dispersion for RNA-Seq data with DESeq2. Genome Biology, 15:550, 12 2014.

[4] Vân Anh Huynh-Thu, Alexandre Irrthum, Louis Wehenkel, and Pierre Geurts. Inferring regulatory networks from expression data using tree-based methods. PloS one, 5, 09 2010.

[5] Macintosh Cornwell, Mahesh Vangala, Len Taing, Zachary Herbert, Johannes Koster, Bo Li, Hanfei Sun, Taiwen Li, Jian Zhang, Xintao Qiu, Matthew Pun, Rinath Jeselsohn, Myles Brown, X Shirley Liu, and Henry W Long. VIPER: Visualization Pipeline for RNA-seq, a Snakemake workflow for efficient and complete RNA-seq analysis. BMC Bioinf, 19, 04 2018.

[6] Denis Torre, Alexander Lachmann, and Avi Ma’ayan. BioJupies: automated generation of interactive notebooks for RNA-Seq data analysis in the cloud. Cell Systems, 7(5):556–561.e3, 2018.

[7] John J Cole, Bekir A Faydaci, David McGuinness, Robin Shaw, Rose A Maciewicz, Robertson Neil A, and Carl S Goodyear. Searchlight: automated bulk RNA-seq exploration and visualisation using dynamically generated R scripts. BMC Bioinf, 22, 08 2021.

[8] Pengyi Yang, Sean J Humphrey, Senthilkumar Cinghu, Rajneesh Pathania, Andrew J Oldfield, Dhirendra Kumar, Dinuka Perera, Jean YH Yang, David E James, Matthias Mann, and Raja Jothi. Multi-omic profiling reveals dynamics of the phased progression of pluripotency. Cell Systems, 8(5):427–445.e10, 2019.

[9] Li Fang, Yunjin Li, Lu Ma, Qiyue Xu, Fei Tan, and Geng Chen. GRNdb: decoding the gene regulatory networks in diverse human and mouse conditions. Nucleic Acids Research, 49(D1):D97–D103, 11 2020.

[10] Daniel Hurley, Hiromitsu Araki, Yoshinori Tamada, Ben Dunmore, Deborah Sanders, Sally Humphreys, Muna Affara, Seiya Imoto, Kaori Yasuda, Yuki Tomiyasu, Kosuke Tashiro, Christopher Savoie, Vicky Cho, Stephen Smith, Satoru Kuhara, Satoru Miyano, D. Stephen Charnock-Jones, Edmund J. Crampin, and Cristin G. Print. Gene network inference and visualization tools for biologists: application to new human transcriptome datasets. Nucleic Acids Research, 40(6):2377–2398, 11 2011.

[11] Paul Shannon, Andrew Markiel, Owen Ozier, Nitin Baliga, Jonathan Wang, Daniel Ramage, Nada Amin, Benno Schwikowski, and Trey Ideker. Cytoscape: A software environment for integrated models of biomolecular interaction networks. Genome research, 13:2498–504, 12 2003.

[12] Hung Nguyen, Duc Tran, Bang Tran, Bahadir Pehlivan, and Tin Nguyen. A comprehensive survey of regulatory network inference methods using single cell RNA sequencing data. Briefings in Bioinformatics, 22(3), 09 2020.

[13] Ilias Moutsopoulos, Lukas Maischak, Elze Lauzikaite, Sergio A Vasquez Urbina, Eleanor C Williams, Hajk-Georg Drost, and Irina I Mohorianu. noisyR: enhancing biological signal in sequencing datasets by characterizing random technical noise. NAR, 49(14):e83–e83, 06 2021.

[14] Minzhe Zhang, Qiwei Li, Donghyeon Yu, Bo Yao, Wei Guo, Yang Xie, and Guanghua Xiao. Geneck: A web server for gene network construction and visualization. BMC Bioinformatics, 20, 01 2019.

[15] Sara Lopez-Gomollon, Irina I Mohorianu, Gyorgy Szittya, Vincent Moulton, and Tamas Dalmay. Diverse correlation patterns between micrornas and their targets during tomato fruit development indicates different modes of microrna actions. Planta, 236, 08 2012.

[16] Yang Liao, Gordon K Smyth, and Wei Shi. featureCounts: an efficient general purpose program for assigning sequence reads to genomic features. Bioinformatics, 30(7):923–930, 11 2013.

[17] Marie-Agnes Dillies, Andrea Rau, Julie Aubert, Christelle Hennequet-Antier, Marine Jeanmougin, Nicolas Servant, Celine Keime, Guillemette Marot, David Castel, Jordi Estelle, Gregory Guernec, Bernd Jagla, Luc Jouneau, Denis Laloe, Caroline Le Gall, Brigitte Schaeffer, Stephane Le Crom, Mickael Guedj, Florence Jaffrezic, and French StatOmique Consortium. A comprehensive evaluation of normalization methods for Illumina high-throughput RNA sequencing data analysis. Briefings in Bioinformatics, 14, 09 2012.

[18] B Bolstad, RA Irizarry, Magnus Åstrand, and T Speed. A comparison of normalization methods for high density oligonucleotide array data based on bias and variance. Bioinformatics, 19:185–93, 02 2003.

[19] Ali Mortazavi, Brian Williams, Kenneth Mccue, Lorian Schaeffer, and Barbara Wold. Mapping and quantifying mammalian transcriptomes by rna-seq. Nature methods, 5:621–8, 08 2008.

[20] Uku Raudvere, Liis Kolberg, Ivan Kuzmin, Tambet Arak, Priit Adler, Hedi Peterson, and Jaak Vilo. g:Profiler: a web server for functional enrichment analysis and conversions of gene lists (2019 update). NAR, 47(W1):W191–W198, 05 2019.

[21] Hsi-Yuan Huang, Yang-Chi-Dung Lin, Jing Li, Kai-Yao Huang, Sirjana Shrestha, Hsiao-Chin Hong, Yun Tang, Yi-Gang Chen, Chen-Nan Jin, Yuan Yu, Jia-Tong Xu, Yue-Ming Li, Xiao-Xuan Cai, Zhen-Yu Zhou, Xiao-Hang Chen, Yuan-Yuan Pei, Liang Hu, Jin-Jiang Su, Shi-Dong Cui, Fei Wang, Yue-Yang Xie, Si-Yuan Ding, Meng-Fan Luo, Chih-Hung Chou, Nai-Wen Chang, Kai-Wen Chen, Yu-Hsiang Cheng, Xin-Hong Wan, Wen-Lian Hsu, Tzong-Yi Lee, Feng-Xiang Wei, and Hsien-Da Huang. miRTarBase 2020: updates to the experimentally validated microRNA–target interaction database. Nucleic Acids Research, 48(D1):D148–D154, 2019.

[22] V. Matys, O. V. Kel-Margoulis, E. Fricke, I. Liebich, S. Land, A. Barre-Dirrie, I. Reuter, D. Chekmenev, M. Krull, K. Hornischer, N. Voss, P. Stegmaier, B. Lewicki-Potapov, H. Saxel, A. E. Kel, and E. Wingender. TRANSFAC® and its module TRANSCompel®: transcriptional gene regulation in eukaryotes. Nucleic Acids Research, 34(suppl 1):D108–D110, 01 2006.

[23] Yumei Li, Xinzhou Ge, Fanglue Peng, Wei Li, and Jingyi Li. Exaggerated false positives by popular differential expression methods when analyzing human population samples. Genome Biology, 23, 12 2022.

[24] Philip Ewels, Mans Magnusson, Sverker Lundin, and Max Kaller. MultiQC: summarize analysis results for multiple tools and samples in a single report. Bioinformatics, 32(19):3047–3048, 06 2016.

[25] Alexander Dobin, Carrie A Davis, Felix Schlesinger, Jorg Drenkow, Chris Zaleski, Sonali Jha, Philippe Batut, Mark Chaisson, and Thomas R Gingeras. STAR: ultrafast universal RNA-seq aligner. Bioinformatics, 29(1):15–21, 10 2012.

[26] Ben Langmead and Steven L Salzberg. Fast gapped-read alignment with bowtie 2. Nature methods, 9:357–9, 03 2012.

[27] Yong Zhang, Tao Liu, Clifford A Meyer, Jerome Eeckhoute, David S Johnson, Bradley E Bernstein, Chad Nusbaum, Richard M Myers, Myles Brown, Wei Li, and X Shirley Liu. Model-based analysis of ChIP-seq (MACS). Genome Biology, 9:R137, 10 2008.

[28] Fidel Ramírez, Devon P Ryan, Björn Grüning, Vivek Bhardwaj, Fabian Kilpert, Andreas S Richter, Steffen Heyne, Friederike Dündar, and Thomas Manke. deepTools2: a next generation web server for deep-sequencing data analysis. Nucleic Acids Research, 44(W1):W160–W165, 04 2016.

[29] Zhimin Li, Yu Sun, Maolin He, and Jianwei Liu. Differentially-expressed mrnas, micrornas and long noncoding rnas in intervertebral disc degeneration identified by rna-sequencing. Bioengineered, 12(1):1026–1039, 2021.

[30] Ana Kozomara, Maria Birgaoanu, and Sam Griffiths-Jones. miRBase: from microRNA sequences to function. Nucleic Acids Research, 47(D1):D155–D162, 2018.

[31] Kay Prüfer, Udo Stenzel, Michael Dannemann, Richard E Green, Michael Lachmann, and Janet Kelso. PatMaN: Rapid alignment of short sequences to large databases. Bioinformatics, 24:1530–1, 2008.

[32] Jake R Conway, Alexander Lex, and Nils Gehlenborg. UpSetR: an R package for the visualization of intersecting sets and their properties. Bioinformatics, 33(18):2938–2940, 06 2017.

