## Supplementary figures for "bulkAnalyseR: An accessible, interactive pipeline for analysing and sharing bulk multi-modal sequencing data"

### 1 Workflow Diagram

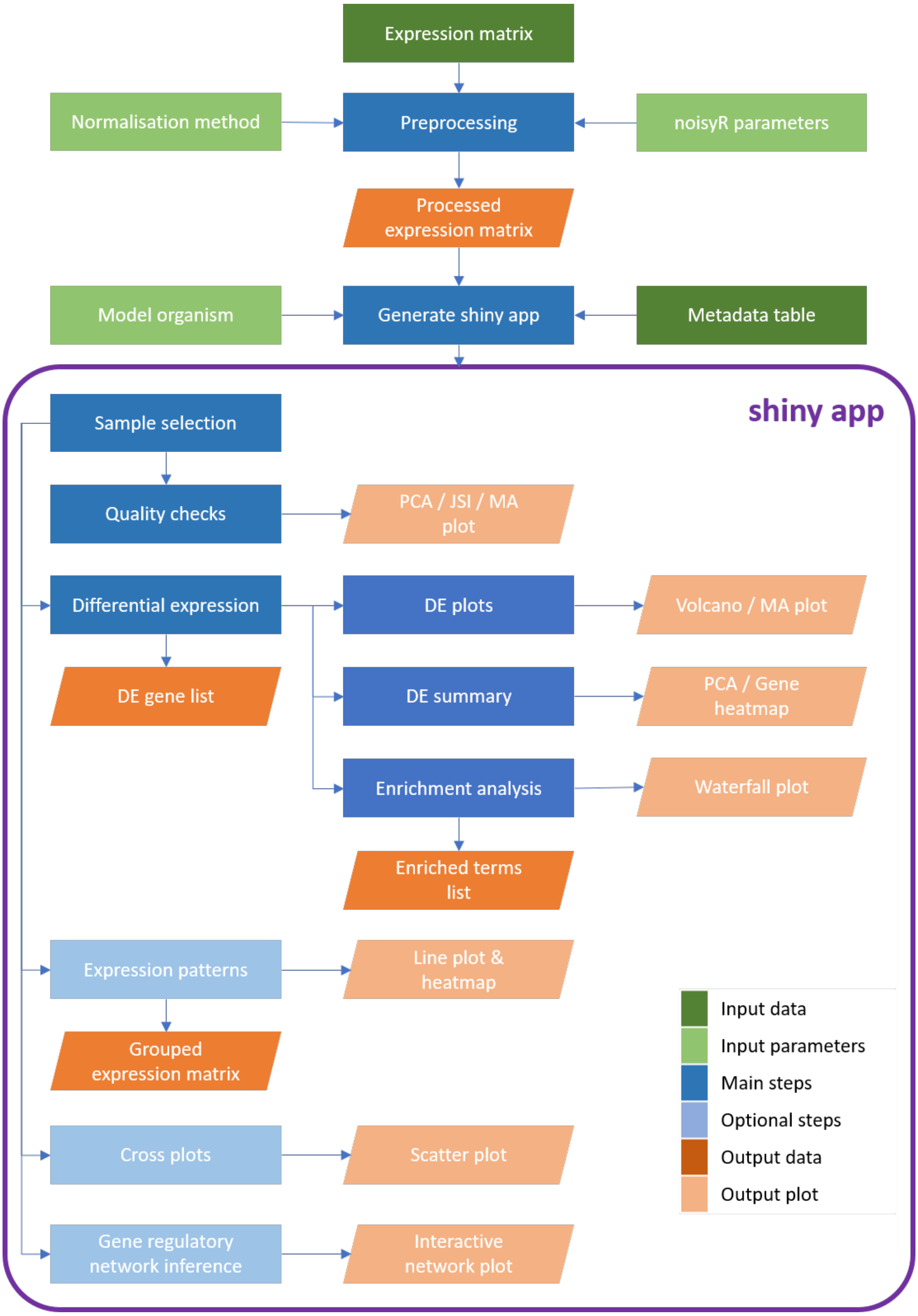


Figure S 1: **Workflow diagram illustrating the bulkAnalyseR pipeline.** The input comprises a processed (i.e. normalised, noise corrected) expression matrix, and a metadata table. Using **bulkAnalyseR**, all standard steps related to differential expression analyses are handled seamlessly e.g. DE call, pairwise comparison of differential expression outputs using cross plots and upset plots, and the inference of localised Gene Regulatory Network.

### 2 Pre-processing Overview

A

**Algorithm 1:** bulkAnalyseR preprocessing

**input** : expression.matrix the expression matrix; rows correspond to genes and columns correspond to samples
**parameters:** *output.plot* (whether to create an expression-similarity plot for the noise analysis; default is FALSE)
*normalisation.method* (the normalisation method to be used, quantile, RPM, TMM or DESeq2; default is quantile)

1 Perform noisyR denoising (count version) using any additional parameters provided
2 If *output.plot*=TRUE, noisyR expression similarity plot is output to console
3 Perform normalisation using *normalisation.method*

**output** : The denoised, normalised expression matrix; some rows (genes) may have been removed by noisyR.

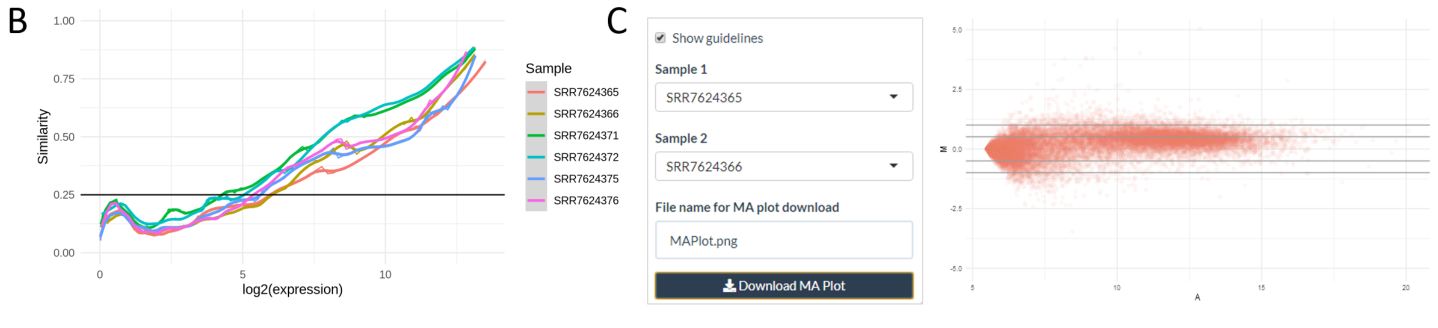


Figure S 2:
A Pseudocode summarising the pre-processing performed in `preprocessExpressionMatrix`.
B Line plot output from the noisyR noise identification pipeline showing the Pearson Correlation Coefficient (y-axis), calculated on windows of increasing average abundance (x-axis, $log_2$ scale), using count matrix-based noise removal approach. This case study illustrates the results for the 0h, 12h and 36h samples.
C MA plot comparing expression levels for 0h rep 1 and rep 2 samples from the QC panel. This comparison can be performed between any two samples.

### 3 Differential expression comparison

Discrepancies between frequently used pipelines for inferring differential expression are often observed but rarely tackled, with researchers usually choosing either one set of results per experiment, or focusing on the the union/ intersection of results across methods. A recent study by Li et al highlighted the variability deriving the False Discovery Rate control, leading to inconsistent results across pipelines. These differences can be reduced significantly by controlling the (technical) noise, as underlined also in the noisyR study. For the Yang et al case study, we highlight these differences by contrasting the DESeq2 and edgeR DE outputs, when the pipelines are run on the same inputs (noisy and denoised matrices are assessed, alongside different normalisation methods). The results are summarised using UpSet plots, and support the noise correction step.

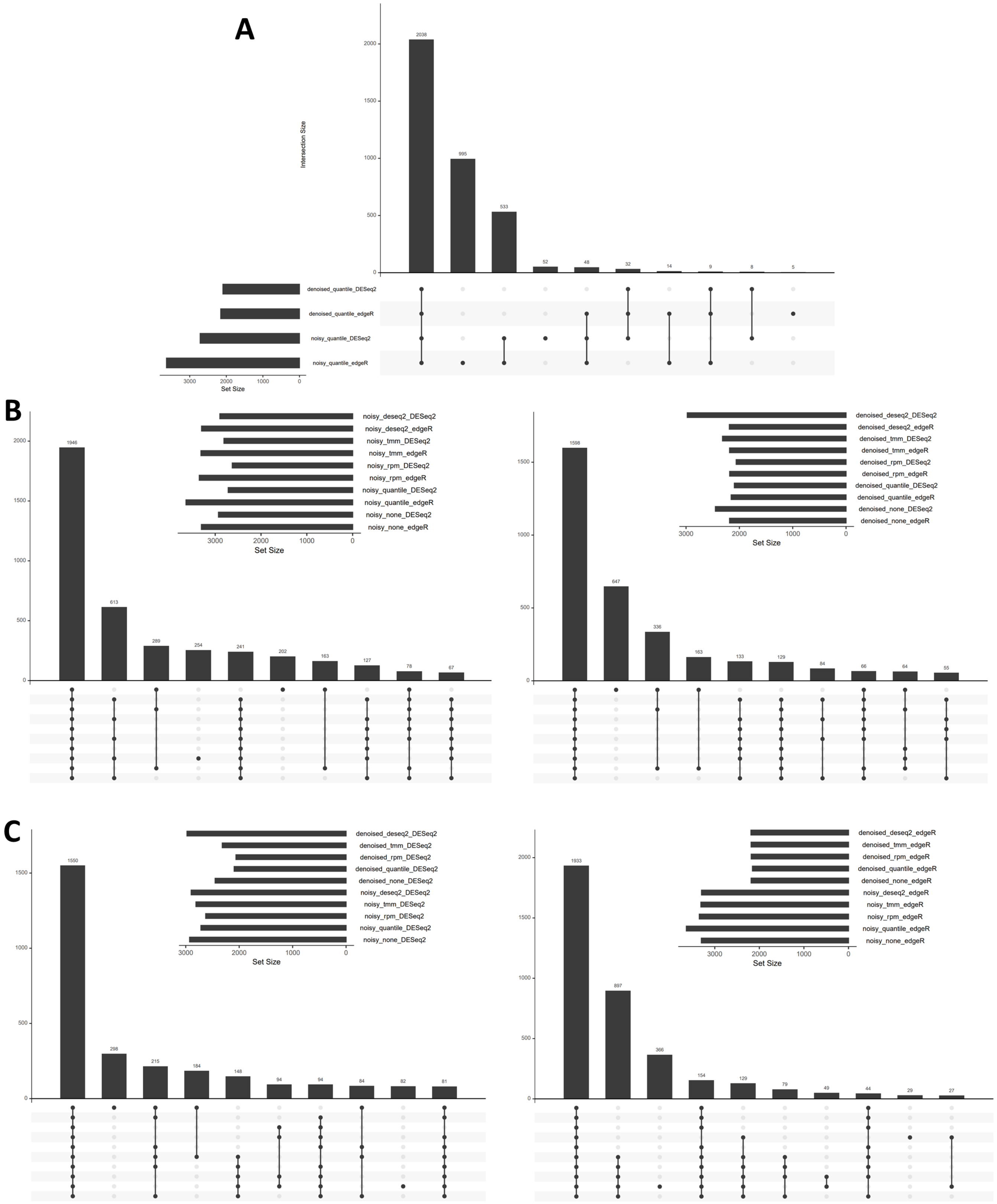


Figure S 3: **Comparison of DE results across pipelines and pre-processing methods**. The largest discrepancies between DE calls are observed when noisy input data is used; the variability is reduced after denoising.
A UpSet plot summarising DE sets with or without the noise correction, for a DE call performed using DESeq2 or edgeR; the expression levels were normalised using quantile normalisation.
B UpSet plot summarising DE sets (without the noise correction, left panel; denoised, right panel) for a DE call performed using DESeq2 or edgeR and various normalisation approaches (DESeq2, TMM, RPM, quantile).
C UpSet plot summarising DE sets with or without the noise correction (the analysis was performed using DESeq2, left panel; the analysis was performed using edgeR, right panel) using different normalisation approaches (DESeq2, TMM, RPM, quantile).

### 4 App Panels and Functionality [mRNAseq]

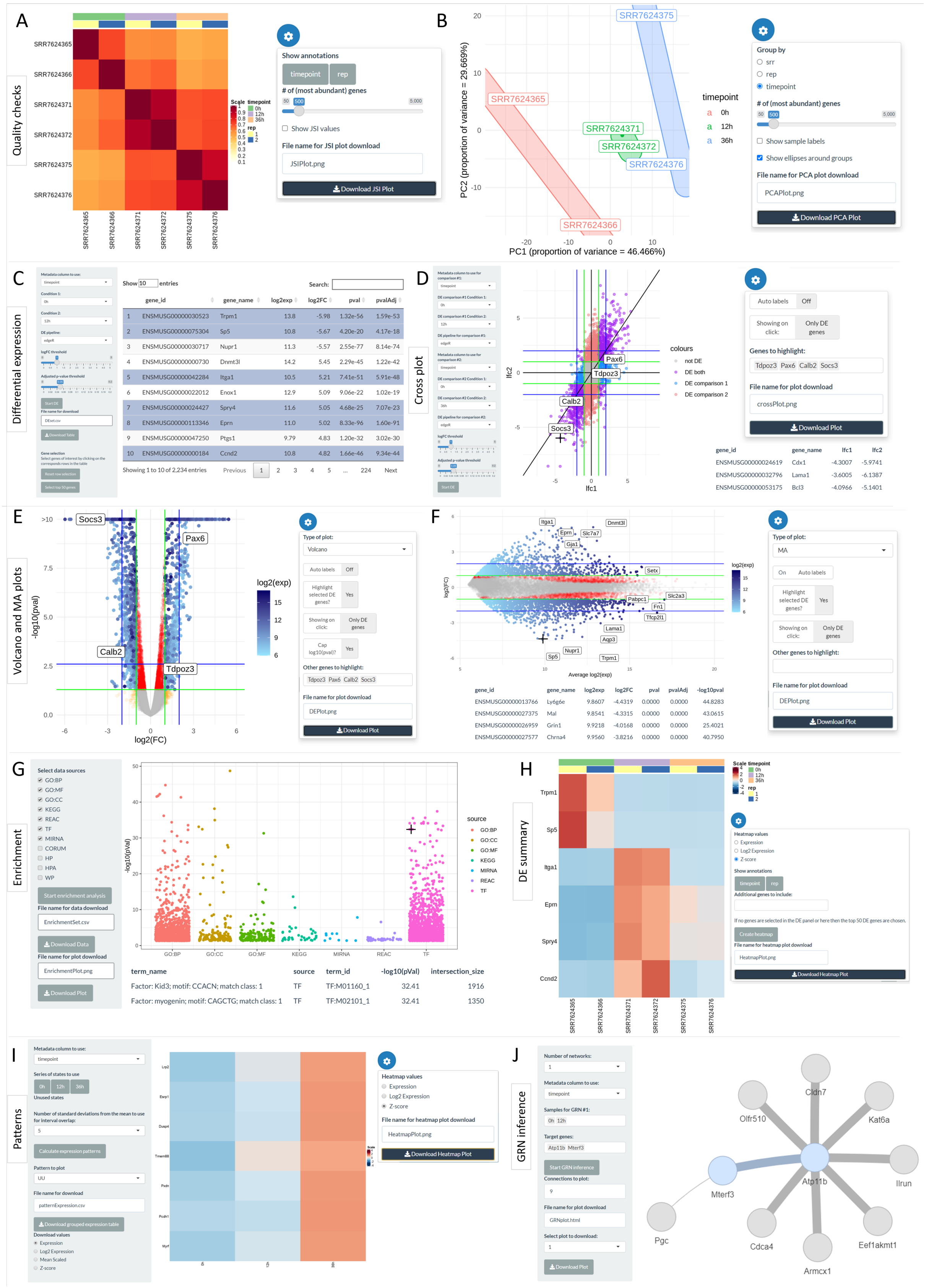


Figure S 4: Visualisation of shiny app functionality

**bulkAnalyseR summary plots on a bulk mRNAseq case study, Yang *et al* data**

***Quality checks***

A Jaccard Similarity Index (JSI) on the 500 most abundant genes, across samples. The JSI can be reordered by metadata columns; the number of most abundant genes is user-defined.
B PCA on the top 500 most abundant genes, coloured by time point. The groups are user-defined, based on any metadata column; labels and ellipses are customisable i.e. can be added or removed.

***Differential expression (DE)***

C Table of top 10 DE genes selected using $log_2$FC threshold 1 and adjusted p-value threshold 0.05 (on a Benjamini-Hochberg multiple testing correction), comparing 0h and 12h from the Yang *et al* dataset.

***Cross plot***

D Scatter (cross) plot comparing $log_2$FC for DE analysis on 0h vs 12h (x-axis) and 0h vs 36h (y-axis). Green guide lines show the $log_2$FC DE threshold and blue guide lines show twice the DE threshold. The colour code indicates the DE status in both comparisons: purple (genes DE in both comparisons), blue (genes DE only for comparison 1), red (genes DE only for comparison 2) and grey (genes DE in neither comparison). A table showing the $log_2$FC values for the closest 4 genes to the click position is displayed below.

***Volcano and MA plots***

E-F Volcano (E) and MA (F) plots comparing 0h and 12h samples. Green guide lines show the $log_2$FC DE threshold and $log_{10}$ of the p-value threshold; blue guide lines indicate thresholds twice as strict. The colour gradient of the DE genes is proportional to $log_2$expression. Labels are shown for (E) selected genes (Socs3, Pax6, Calb2 and Tdpoz3) and (F) auto-selected genes, and tables showing DE results for the closest genes to click position are displayed below.

***Enrichment***

G Scatter plot showing $log_{10}$p-value from g:profiler enrichment for Gene Ontology (GO), KEGG, Reactome (REAC), miRNA and Transfac (TF) terms. Other sources can be included. A table showing the terms and $log_{10}$p values for the closest 2 terms to the click position is shown below.

***DE summary***

H Heatmap showing Z-score transformed expression levels for genes selected in DE panel. If no genes are selected then the top 50 genes by $|log_2\text{FC}|$ are used. Further genes can be added by name. The order of samples can be altered as in panel A.

***Patterns***

I Heatmap showing Z-score transformed mean expression of the genes assigned to the chosen pattern, UU, in each timepoint. Gene names are shown if less than 50 genes are present. The pattern identification is done on confidence intervals, calculated for each gene, per condition; the CI is built on the mean $\pm$ a user-defined multiple of standard deviations. The pattern between two conditions is defined as straight (S) if the CIs overlap and up (U) or down (D) if they do not.

***GRN inference***

J Visualisation of top 9 connections in an inferred GRN (using GENIE3) using 0h and 12h samples; the selected DE genes are Atp11b and Mterf3. The width of the connecting edges is proportional to the weight from the adjacency matrix.

### 5 App Panels and Functionality [ChIPseq]

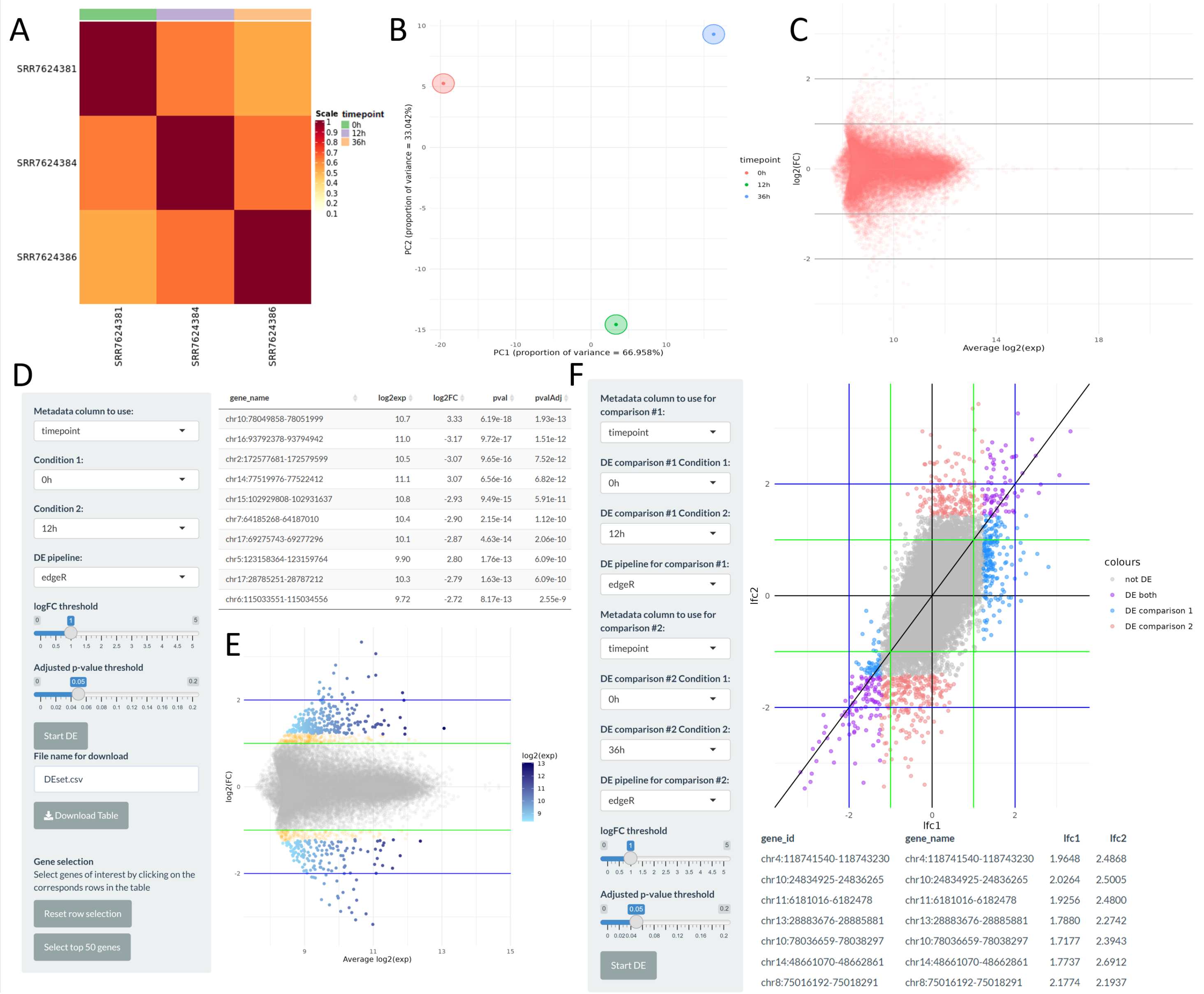


Figure S 5: **bulkAnalyseR summary plots on a bulk ChIPseq case study, Yang *et al* data**
A Jaccard Similarity Index (JSI) on the 500 highest amplitude peaks, across samples. The JSI can be reordered by metadata columns; the number of highest amplitude peaks is user-defined.
B PCA using top 500 highest amplitude peaks, coloured by timepoint. The groups are user-defined, based on any metadata column; labels and ellipses are customisable i.e. can be added or removed.
C MA plot comparing expression levels for 0h and 12h sample from the QC panel. This comparison can be performed between any two samples.
D Table of top 10 DE peaks selected using $log_2$FC threshold 1 and adjusted p-value threshold 0.05 (with Benjamini-Hochberg multiple testing correction), comparing 0h and 12h samples
E MA plot comparing 0h and 12h samples in differential expression. Green guide lines show the $log_2$FC DE threshold and $log_{10}$ of the p-value threshold; blue guide lines indicate thresholds twice as strict. The colour gradient of the DE peaks is proportional to $log_2$expression.
F Scatter (cross) plot comparing $log_2$FC for a DE analysis on 0h vs 12h (x-axis) and 0h vs 36h (y-axis). Guide lines and labels are as in panel E. The colour code indicates the DE status in both comparisons: purple (peaks DE in both comparisons), blue (peaks DE only for comparison 1), red (peaks DE only for comparison 2) and grey (peaks DE in neither comparison). A table showing the $log_2$FC values for the closest 7 peaks to the click position is shown below.

### 6 App Panels and Functionality [sRNAseq/ microRNAs]

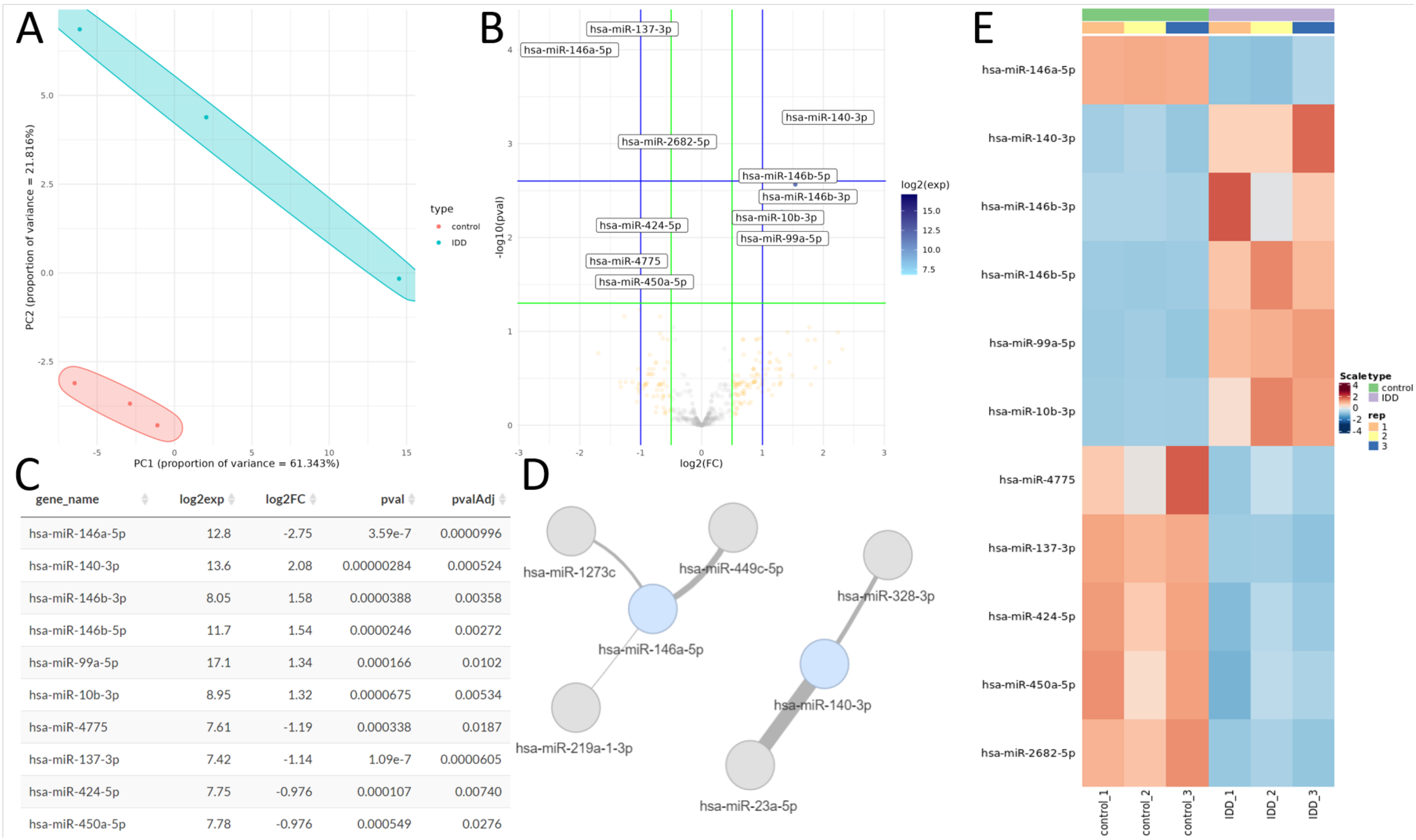


| gene_name | log2exp | log2FC | pval | pvalAdj |
|---|---|---|---|---|
| hsa-miR-146a-5p | 12.8 | -2.75 | 3.59e-7 | 0.0000996 |
| hsa-miR-140-3p | 13.6 | 2.08 | 0.00000284 | 0.000524 |
| hsa-miR-146b-3p | 8.05 | 1.58 | 0.0000388 | 0.00358 |
| hsa-miR-146b-5p | 11.7 | 1.54 | 0.0000246 | 0.00272 |
| hsa-miR-99a-5p | 17.1 | 1.34 | 0.000166 | 0.0102 |
| hsa-miR-10b-3p | 8.95 | 1.32 | 0.0000675 | 0.00534 |
| hsa-miR-4775 | 7.61 | -1.19 | 0.000338 | 0.0187 |
| hsa-miR-137-3p | 7.42 | -1.14 | 1.09e-7 | 0.0000605 |
| hsa-miR-424-5p | 7.75 | -0.976 | 0.000107 | 0.00740 |
| hsa-miR-450a-5p | 7.78 | -0.976 | 0.000549 | 0.0276 |

Figure S 6: **bulkAnalyseR summary plots on a bulk sRNAseq case study (with a focus on mature microRNA expression), Li *et al* data**
A PCA using top 100 highest amplitude miRNAs, coloured by type.
B Volcano plot illustrating the control vs IDD differential expression.
C Table of top 10 DE miRNAs selected using $log_2$FC threshold 0.5 and adjusted p-value threshold 0.05 (with Benjamini-Hochberg multiple testing correction), comparing control and IDD samples
D Inferred GRN on miRNA data, from targets hsa-miR-146a-5p and hsa-miR-140-3p.
E Heatmap showing Z-score transformed expression levels for miRNAs selected in DE panel.

### 7 Online Deployment

Apps created with **bulkAnalyseR** are standalone and can be easily deployed online. Publication on user-owned public websites is ideal for sharing data and encouraging other researchers to reproduce an extend analyses and results.
An easy, universally available, option for hosting shiny apps is the shinyapps.io platform. To deploy an app, the user needs to create an ACCOUNT and employ the dedicated *rsconnect* package available on the R platform (RStudio). The app will then be deployed at https://[ACCOUNT].shinyapps.io/shinyApp/. Sample code for this task is presented below. For more detailed guidance please refer to the shinyapps.io user guide.

```
install.packages("rsconnect")
rsconnect::setAccountInfo(
    name = "<ACCOUNT>",
    token = "<TOKEN>",
    secret = "<SECRET>"
) # these can be found in your account's Profile -> Tokens page
rsconnect::deployApp("shinyApp/")
```
